# Single-Molecule Imaging Reveals the Mechanism of Bidirectional Replication Initiation in Metazoa

**DOI:** 10.1101/2024.03.28.587265

**Authors:** Riki Terui, Scott Berger, Larissa Sambel, Dan Song, Gheorghe Chistol

## Abstract

Metazoan genomes are copied bidirectionally from thousands of replication origins. Replication initiation entails the assembly and activation of two CMG (Cdc45•Mcm2-7•GINS) helicases at each origin. This requires several firing factors (including TopBP1, RecQL4, DONSON) whose exact roles remain unclear. How two helicases are correctly assembled and activated at every single origin is a long-standing question. By visualizing the recruitment of GINS, Cdc45, TopBP1, RecQL4, and DONSON in real time, we uncovered a surprisingly dynamic picture of initiation. Firing factors transiently bind origins but do not travel with replisomes. Two Cdc45 simultaneously arrive at each origin and two GINS are recruited together, even though neither protein can dimerize. The synchronized delivery of two GINS is mediated by DONSON, which acts as a dimerization scaffold. We show that RecQL4 promotes DONSON dissociation and facilitates helicase activation. The high fidelity of bidirectional origin firing can be explained by a Hopfield-style kinetic proofreading mechanism.

## Introduction

To duplicate their large genomes in a timely fashion, eukaryotes initiate DNA replication in parallel from thousands of sites called origins. Importantly, DNA is replicated bi-directionally from each origin, ensuring that each chromosome is fully replicated regardless of the exact number or location of origins. How bidirectional replication is faithfully established at each origin is a long-standing question in the field.

Eukaryotic replication occurs in a few highly regulated stages (Figure 1A). During G1 chromatin is “licensed” for replication, wherein two Mcm2-7 (Mini-chromosome maintenance proteins 2-7) complexes are loaded onto double stranded DNA (dsDNA) at each origin. Each dimer of Mcm2-7 forms a stable pre-replication complex (pre-RC), also known as a “dormant origin” ^1,2^. During S-phase, GINS (a complex of Sld5, Psf1, Psf2, and Psf3) and Cdc45 (Cell division cycle 45) are recruited to the dormant origin to assemble two Cdc45•Mcm2-7•GINS (CMG) helicases. Next, the pair of helicases is activated, and each CMG begins to unwind DNA ^3^.

**Figure 1:**
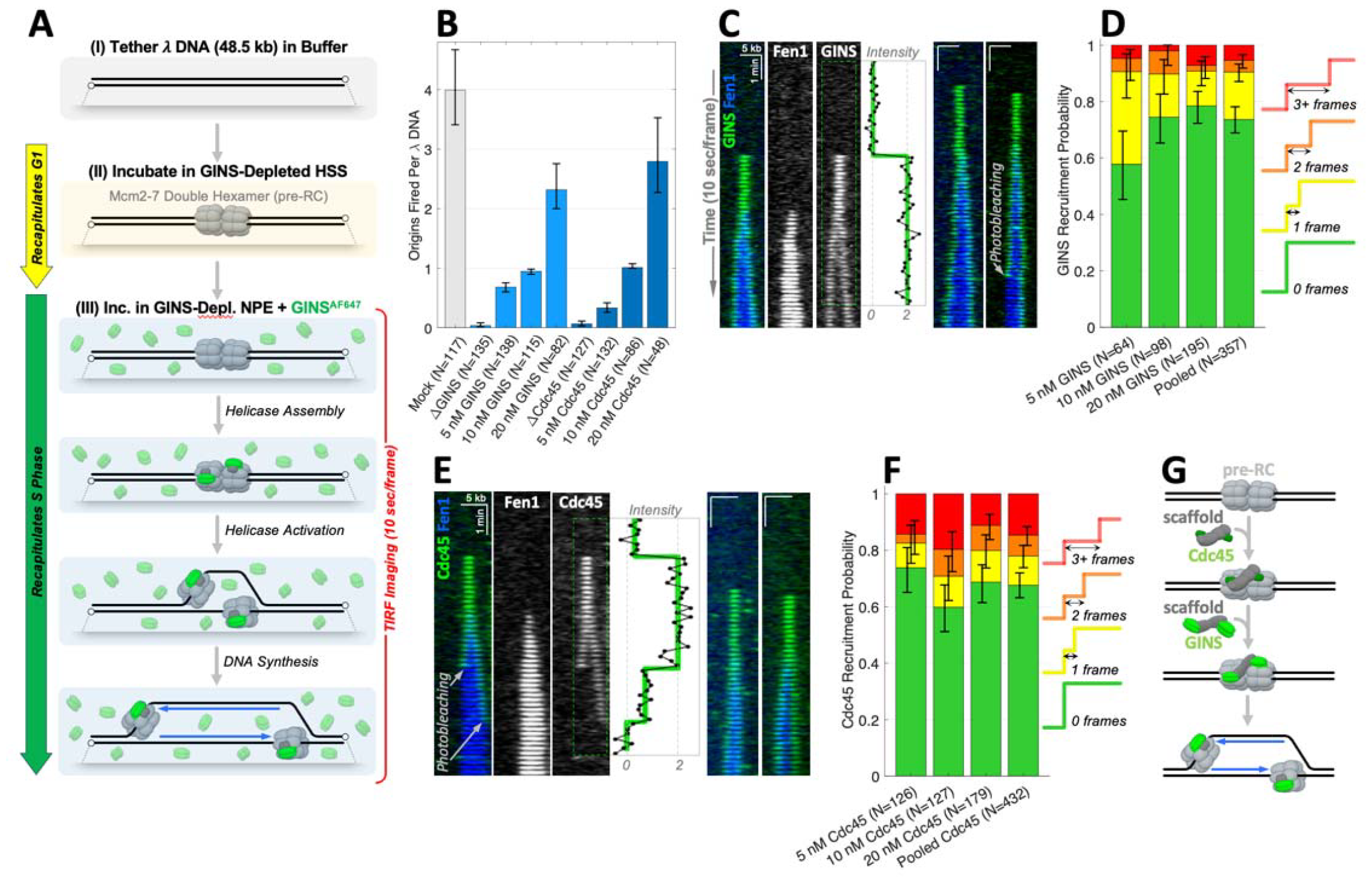
A Dimer of GINS and a Dimer of Cdc45 Are Recruited to Each Mcm2-7 Double Hexamer. **(A)** Workflow of single-molecule experiment to visualize GINS recruitment during replication initiation. GINS^AF647^ is shown in green, nascent DNA labeled with Fen1^mKikGR^ is shown in blue. Firing factors and other replication proteins are not shown for clarity. **(B)** Origin firing efficiency for GINS^AF647^ (light blue) and Cdc45^AF647^ (dark blue) titration experiments compared to mock-depleted (gray), GINS-depleted, and Cdc45-depleted reactions (N – number of DNA molecules for each condition). Origins were counted u ing the Fen1^mKikGR^ channel. **(C)** Representative kymograms showing the recruitment of GINS^AF647^ (green) during initiation. Fen1^mKikGR^ (blue) serves as a marker of nascent DNA. Inset: integrated GINS^AF647^ signal (black) and a changepoint fit to the raw signal (green). An ensemble assay was used to verify that the recombina t GINS was biochemically active (Figure S1C-D). As shown previously, Alexa Fluor 647 does not interfere with GINS function ^22^. **(D)** Breakdown of how two copies of GINS were recruited to the origin. At most origins, two GINS molecules appear simultaneously (0 frames, green). At the remaining origins, two copies of GINS show up sequentially, where the second molecule appears 1 frame (yellow), 2 frames (orange), or 3+ frames (r d) after the first GINS (N – number of fired origins where both copies of GINS were fluorescently labeled). **(E)** Representative kymograms showing the recruitment of Cdc45^AF647^ (green) during initiation. Inset: integrated Cdc45^AF647^ signal (black) and the changepoint fit to the raw signal (green). A biochemical assay was used to verify that fluorescent labeling did not impair Cdc45’s function (Figure S1E-G). **(F)** Breakdown of how two copies of Cdc45 were recruited to the origin (N – number of fired origins where both copies of Cdc45 were labeled). **(G)** Model illustrating how two copies of GINS or Cdc45 could be simultaneously recruited to the pre-RC. Cdc45 recruitment is depicted before GINS binding because previous studies have shown that Cdc45 is recruited to chromatin in the absence of GINS, but not vice-versa ^40,43^. In all bar plots, error-bars represent the 95% confidence interval (CI) for the mean estimated via bootstrapping.

Several other replication proteins (polymerases, processivity clamps, nucleases, structural scaffolds, etc.) are recruited to each helicase, forming a mature replisome. Helicase assembly and activation involves several “firing factors” (TopBP1, RecQL4, DONSON, Treslin-MTBP, Mcm10) whose exact functions are poorly understood ^4^.

TopBP1 (Topoisomerase II Binding Protein 1) is a large protein consisting of several phosphopeptide-binding domains BRCT (BRCA1 C-terminal) interspersed with disordered regions. TopBP1 is required for recruiting Cdc45 and GINS to chromatin, but its mechanism of action is unclear ^5,6^. In addition to its role in initiation, TopBP1 is involved in the DNA damage response as an allosteric activator of the ATR (Ataxia Telangiectasia and Rad53 related) kinase ^7^. TopBP1 oligomerization is critical for its function as an ATR activator ^8^, but it is unclear how TopBP1 oligomerization regulates origin firing.

RecQL4 (RecQ Like helicase 4) contains a domain distantly related to the yeast Sld2 – an essential firing factor in yeast. RecQL4 also contains an ATPase domain and long disordered regions. There are conflicting reports about RecQL4’s role in replication initiation: it is thought to be involved in helicase assembly or CMG activation ^9–12^. RecQL4 was also implicated in other aspects of genome maintenance, making it especially challenging to interrogate its role in initiation ^13^. Finally, it is unclear if RecQL4 travels with the replisome, and how that modulates its function.

DONSON was recently identified as a novel regulator of metazoan replication initiation with no direct homolog in yeast. The emerging consensus is that DONSON facilitates GINS recruitment to the Mcm2-7 double hexamer ^14–17^. DONSON was also implicated in DNA damage response as a fork protection factor ^18,19^. How it accomplishes these two distinct functions remains unclear. Although DONSON has been reported to travel with the replisome, its function at the replication fork remains unknown ^15,18^.

Replication initiation is also regulated by kinases Cdk2/CDK and Cdc7/DDK, which phosphorylate firing factors and Mcm2-7 ^4,20^. In yeast, CDK is thought to promote the formation of a pre-loading complex (pre-LC) consisting of Dpb11 (the yeast homolog of TopBP1), Sld2, GINS, and Polε (the leading strand polymerase). According to the yeast paradigm, the pre-LC is recruited to dormant origins during initiation ^21^. A similar model was recently proposed for metazoa, with the putative pre-LC consisting of either TopBP1•DONSON•GINS•Polε or DONSON•GINS•Cdc45•Polε ^16,17^. However, there is no direct evidence showing that these proteins are recruited to DNA as complex.

Origin firing remains the most poorly understood aspect of replication biology. Specifically, how are two CMG helicases correctly assembled and activated at thousands of origins in each dividing cell? What is the function of each firing factor, and how are they coordinated?

To systematically dissect the mechanism of replication initiation, we employed single-molecule imaging to directly monitor the recruitment of GINS, Cdc45, TopBP1, RecQL4, and DONSON to origins. We found that two copies of GINS are simultaneously recruited to the pre-RC. This is facilitated by DONSON, which acts as a dimerization scaffold for GINS. Similarly, we found that two copies of Cdc45 are also simultaneously recruited to the pre-RC, suggesting that a yet-unknown protein acts as a dimerization scaffold for Cdc45. Single-molecule imaging revealed that TopBP1, RecQL4, and DONSON transiently dock the pre-RC before origin firing, but none of them travel with replication forks. Our observations do not support the pre-loading complex model. Instead, our data indicate that replication initiation is a highly dynamic process that entails the stepwise assembly of several short-lived intermediates. We propose that these metastable states facilitate kinetic proofreading that enforces the two-fold symmetry required for bi-directional replication.

## RESULTS

### A Single-Molecule Assay to Visualize Replication Initiation in Real Time

To directly visualize replication initiation, we adapted a single-molecule imaging approach called KEHRMIT (<u>K</u>inetics of the <u>E</u>ukaryotic <u>H</u>elicase by <u>R</u>eal-time <u>M</u>olecular Imaging and <u>T</u>racking) ^22^. First, λ DNA was flow-stretched and immobilized in a microfluidic flow cell (Figure 1AI). Next, DNA was incubated with High-Speed Supernatant (HSS) of cytoplasmic extract from *Xenopus laevis* eggs (Figure 1AII). HSS supports efficient loading of Mcm2-7 onto DNA without initiating replication. Since metazoan origins are not defined by conserved DNA sequences, pre-RCs are loaded at random locations on the λ substrate. Licensed DNA was incubated with Nucleo-Plasmic Extract (NPE), which supports efficient DNA replication and recapitulates S-phase (Figure 1AIII) ^23^. GINS-depleted NPE was supplemented with GINS^AF647^ (GINS labeled with Alexa Fluor 647) and Fen1^mKikGR^ (a marker of DNA synthesis ^24^). The replication reaction was monitored using a Total Internal Reflection Fluorescence (TIRF) microscope ^25^.

In the original version of KEHRMIT, origins were fired for only a few minutes in the presence of high concentrations of GINS^AF647^ (∼200 nM), which was incompatible with single-molecule imaging. Excess GINS^AF647^ was flushed from the flow cell and fluorescently labeled helicases were imaged in GINS-depleted extract. In the current study, lower concentrations (5-30 nM) of GINS^AF647^ were present in the reaction for the entire experiment, enabling us to visualize GINS recruitment to origins in real time.

### A Dimer of GINS and a Dimer of Cdc45 Are Recruited to Each Mcm2-7 Double Hexamer

It has long been known that DNA is replicated bidirectionally from each origin of replication ^26^. How bidirectionality is established and enforced during origin firing is not known. To address this question, we visualized the events upstream of origin firing – namely GINS and Cdc45 recruitment.

First, we imaged origin firing in reactions containing 5-20 nM of GINS^AF647^ (Figure 1B, light blue bars). Shortly after binding to the pre-RC, two GINS^AF647^ molecules began moving bidirectionally away from the origin, indicating that they were successfully incorporated into active helicases (Figure 1C, green signal). Each initiation event was also clearly detected in the Fen1 channel as a growing replication bubble (Figure 1C, blue signal). Intriguingly, two GINS molecules were recruited within less than 10 seconds – the time resolution of our experiment (Figure 1C, inset). This was true for most origin firing events at different GINS^AF647^ concentrations (Figure 1D).

Next, we asked how Cdc45 is recruited during initiation. To this end, we monitored Cdc45^AF647^ binding to the pre-RC using the same workflow as for GINS. In most cases, two copies of Cdc45^AF647^ appeared to be simultaneously recruited to the pre-RC (Figure 1E, inset), regardless of Cdc45 concentration (Figure 1F). The observation that both GINS and Cdc45 are recruited to origins in pairs was surprising because neither protein dimerizes *in vitro* (Figure S1A-B).

We initially hypothesized that two molecules of Cdc45 were recruited sequentially but in rapid succession, making their binding appears simultaneous. If so, the recruitment of the 2^nd^ molecule should be resolvable when Cdc45 binding is rate-limiting. Lowering Cdc45^AF647^ concentration from 20 nM to 5 nM dramatically reduced the probability of origin firing (Figure 1B, dark blue bars), suggesting that in this regime Cdc45 binding was rate-limiting for initiation. However, this did not significantly change the likelihood of simultaneous Cdc45 recruitment (Figure 1F). Similarly, making GINS^AF647^ rate-limiting for origin firing did not significantly change the probability that two GINS were recruited simultaneously (Figure 1D). Although we cannot strictly rule out that Cdc45/GINS recruitment is sequential but highly cooperative, the data suggest that these helicase subunits are loaded as pre-formed dimers.

Since neither GINS nor Cdc45 alone can dimerize, we hypothesized the existence of dimerization scaffolds for GINS and Cdc45 (Figure 1G). Due to structural differences and their unique placement within the helicase, GINS and Cdc45 probably require unique scaffolds. These putative scaffolds should satisfy a few key requirements: they likely oligomerize, they must directly bind their client protein (GINS or Cdc45), and they must bind the pre-RC or another protein that docks to the pre-RC. Several replication firing factors fit these criteria, including TopBP1 and RecQL4.

### TopBP1 Is Recruited to the Pre-RC Before GINS or Cdc45

We initially regarded TopBP1 as the most promising dimerization scaffold candidate because it can bind both GINS and Cdc45 ^27–29^. Additionally, the yeast ortholog of TopBP1 (Dpb11) was thought to be a subunit of the pre-loading complex (pre-LC) proposed to recruit GINS to dormant origins ^21^. To test this idea, we visualized TopBP1 binding to DNA during replication initiation using the same workflow as for GINS. Since full length TopBP1^AF647^ (Figure S2A-B) formed large assemblies that hindered our ability to analyze and interpret the data (Figure S2C), we prepared a truncated construct (Figure S2D-E) that lacks the ATR activation domain^7,30^ (Figure 2A). The truncated protein retained its origin firing activity (Figure 2B, S2F-H) suggesting it was a good surrogate for full-length TopBP1.

**Figure 2:**
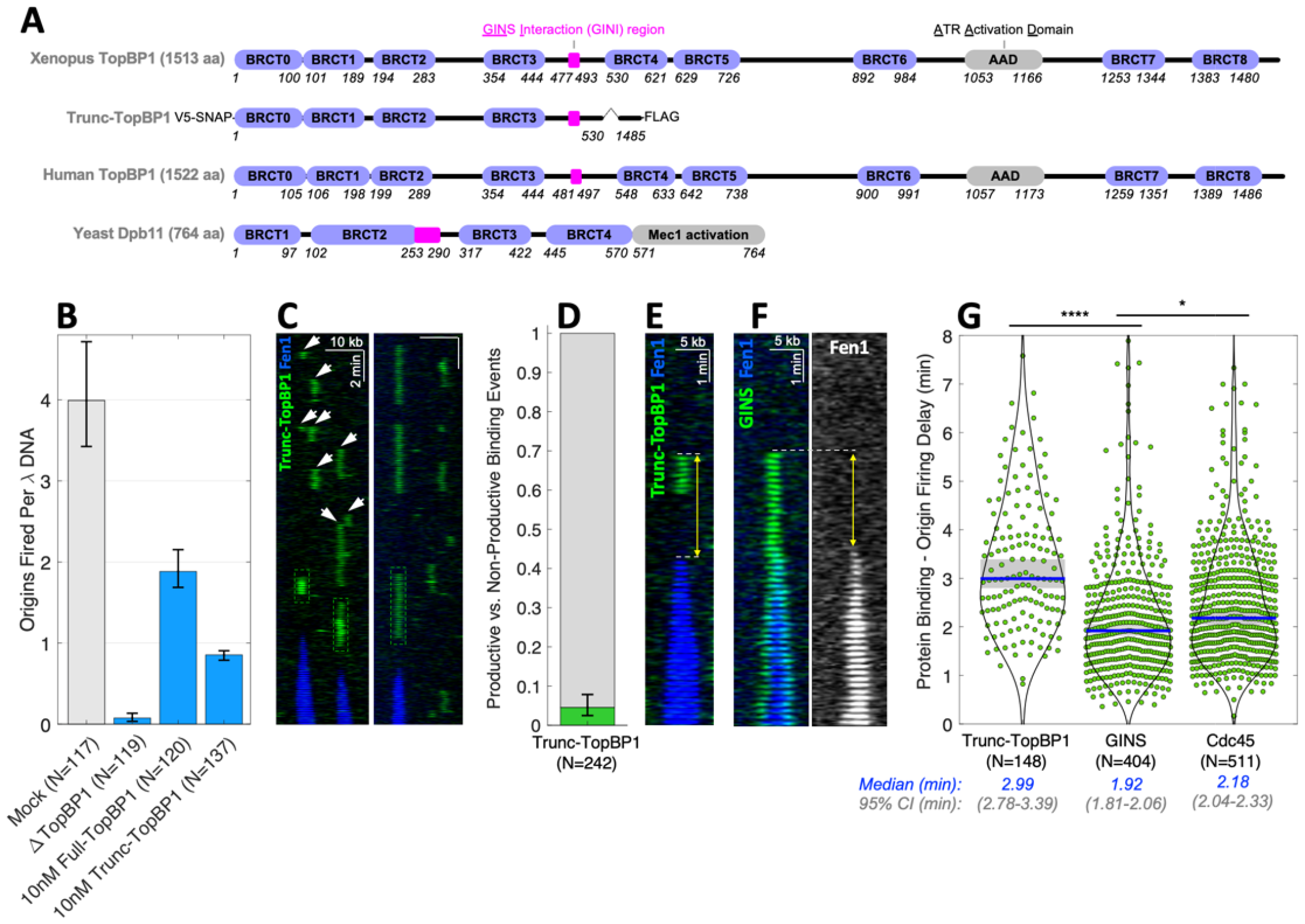
TopBP1 Is Recruited to the Pre-RC Before GINS and Cdc45. **(A)** Domain map of full-length and truncated Xenopus TopBP1 versus human TopBP1 and yeast Dpb11. **(B)** Origin firing efficiency for full-length TopBP1^AF647^ and truncated TopBP1^AF647^ compared to mock-depleted and TopBP1-depleted reactions (N denotes the number of DNA molecules, error bars – 95% CI). Origins were counted using the Fen1^mKikGR^ channel. **(C)** Kymograms showing truncated TopBP1^AF647^ (green) binding during initiation. Dashed boxes mark productive binding events, white arrows - non-productive binding. A biochemical replication assay was used to verify that the SNAP tag used for fluorescent labeling did not impair trunc-TopBP1’s function (Figure S2F-H). Trunc-TopBP1 fluorescent labeling efficiency exceeded 95% (Figure S2E), so the vast majority of trunc-TopBP1 binding events should be visible. **(D)** Probability of productive (green) versus non-productive (gray) trunc-TopBP1^AF647^ binding events (error bars – 95% CI). **(E)** Kymogram illustrating the time delay between trunc-TopBP1^AF647^ binding and origin firing. **(F)** Kymogram illustrating the delay between GINS^AF647^ recruitment and origin firing. **(G)** Distributions of the measured time delay between protein binding and origin firing for trunc-TopBP1^AF647^, GINS^AF647^, and Cdc45^AF647^, each measured in a separate experiment. Blue bar marks the median, gray box - 95% CI for the median. P-values were computed using the two-sample Kolmogorov-Smirnov test. Significance demarcations: ns P>0.05, * P≤0.05, ** P≤0.01, *** P≤0.001, **** P≤0.0001.

TopBP1 recruitment was surprisingly dynamic – it repeatedly bound to and dissociated from DNA, sometimes at the same location (Figure 2C). These transient binding events lasting 1-2 minutes were observed only on licensed DNA (Figure S2I) indicating that TopBP1 is recruited to pre-RCs, consistent with a previous report ^30^. Interestingly, many trunc-TopBP1^AF647^ binding events were non-productive – i.e. did not trigger origin firing (Figure 2C white arrows, Figure 2D). Nevertheless, trunc-TopBP1^AF647^ recruitment was observed shortly before origin firing (Figure 2C, green boxes), suggesting that TopBP1 binding and dissociation are important steps in replication initiation. Importantly, trunc-TopBP1^AF647^ always dissociated before origin firing and did not travel with the replisome.

Simultaneous imaging of TopBP1^AF546^ and GINS^AF647^ (or Cdc45^AF647^) was not possible due to poor origin firing efficiency at the protein concentrations needed for an acceptable signal-to-noise ratio. To test the hypothesis that TopBP1 is the dimerization scaffold for GINS (or Cdc45), we inferred protein recruitment order by comparing the time delay between protein binding and origin firing (Figure 2E-F). On average, trunc-TopBP1^AF647^ bound to DNA ∼3 min before origin firing, approximately 1 minute before either GINS or Cdc45 were recruited (Figure 2G). This finding suggests that TopBP1 is not the dimerization scaffold for GINS or Cdc45, but instead acts upstream of their recruitment.

### RecQL4 Is Recruited to the Pre-RC After GINS

RecQL4 has several properties predicted for the GINS dimerization scaffold. RecQL4 forms stable dimers *in vitro* (Figure S3A-B). RecQL4 is recruited only to licensed chromatin ^10^, suggesting that it binds pre-RCs. Finally, RecQL4 is a distant homolog of the yeast Sld2 (Figure 3A), a subunit of the pre-loading complex hypothesized to recruit GINS to origins.

**Figure 3.**
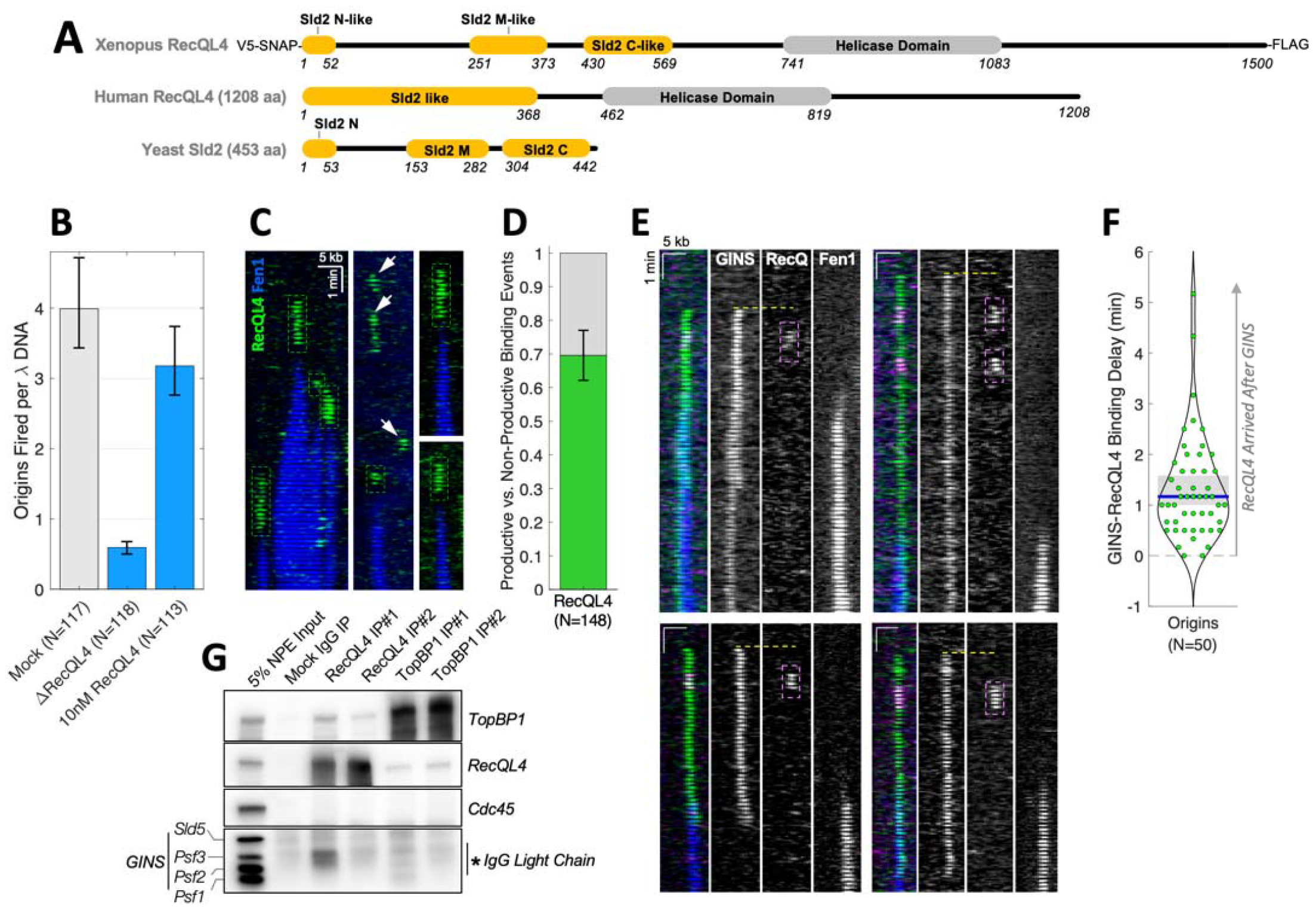
RecQL4 Is Recruited to the Pre-RC After GINS. **(A)** Domain maps of Xenopus and human RecQL4 versus yeast Sld2. **(B)** Origin firing efficiency for RecQL4^AF647^ compared to mock-depleted and RecQL4-depleted reactions (N – number of DNA molecules, error bars – 95% CI). Origins were counted using the Fen1^mKikGR^ channel. **(C)** Kymograms showing the recruitment of RecQL4^AF647^ (green) during replication initiation. Dashed boxes mark productive binding, white arrows - non-productive binding. A biochemical replication assay was used to verify that the SNAP tag did not impair RecQL4’s function (Figure S3C-E). RecQL4 fluorescent labeling efficiency exceeded 90% (Figure S3F), therefore nearly all RecQL4 binding events should be visible. **(D)** Probability of productive (green) versus non-productive (gray) RecQL4^AF647^ binding events (error bars – 95% CI). **(E)** Representative kymograms illustrating the recruitment order of GINS^AF647^ (green) and RecQL4^AF546^ (magenta). In some kymograms, only one GINS molecule is fluorescently labeled. The disappearance of GINS^AF647^ is due to photobleaching. **(F)** Distribution of measured time delays between GINS recruitment and RecQL4 binding for origins where both proteins were detected (blue bar marks the median, gray box – 95% CI for the median). Positive values correspond to RecQL4 arriving after GINS. **(G)** Immunoblots of proteins immunoprecipitated from nucleoplasmic extract using RecQL4 and TopBP1 antibodies raised against two non-overlapping fragments of each protein.

We monitored origin firing in a RecQL4-depleted reaction supplemented with 10 nM of RecQL4^AF647^ (Figure 3B, S3C-F). Like TopBP1, RecQL4’s behavior was very dynamic: it transiently bound to pre-RCs shortly before origin firing (Figure 3C-D, S3G).

To directly test the hypothesis that RecQL4 is a dimerization scaffold for GINS, we simultaneously visualized GINS^AF647^ and RecQL4^AF546^ recruitment during replication initiation. Both GINS and RecQL4 recruitment were clearly detected in 50 origin firing events (Figure 3E), and in all but two of them RecQL4 was recruited after GINS binding (Figure 3F). This finding indicates that RecQL4 is not the GINS dimerization scaffold. Consistent with this conclusion, neither GINS nor Cdc45 co-immunoprecipitated efficiently with RecQL4 (or TopBP1) as would be expected for a dimerization scaffold (Figure 3G). Our data point to RecQL4 acting downstream of GINS recruitment, but before the start of DNA synthesis.

### DONSON and GINS Are Recruited to the Pre-RC at the Same Time

Recently, DONSON was shown to be essential for replication initiation in metazoa ^14–17^. Since DONSON interacts with GINS (Figure 4A-B) ^14–17^ and DONSON can form dimers (Figure S4A-C) ^15,16^, we regarded DONSON as an attractive candidate for the GINS dimerization scaffold.

**Figure 4.**
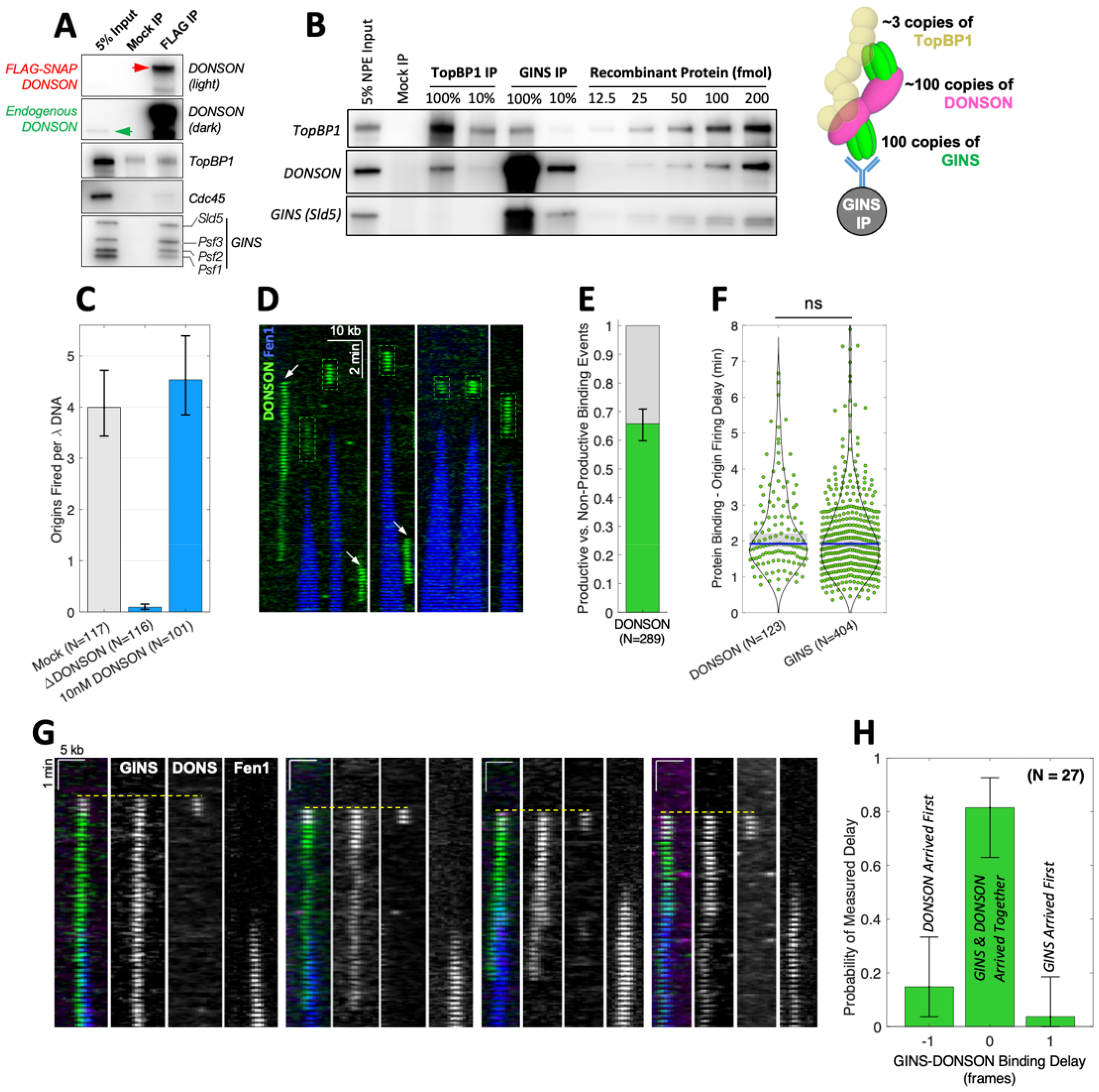
DONSON and GINS Are Simultaneously Recruited to the Pre-RC. **(A)** Immunoblots of proteins immunoprecipitated with FLAG-SNAP-DONSON from NPE. FLAG-SNAP-DONSON migrates slower than the endogenous DONSON from egg extract. **(B)** Quantitative immunoblots of proteins immunoprecipitated from NPE with TopBP1 or GINS versus known amounts of recombinant protein. The cartoon illustrates the approximate co-IP stoichiometry for the GINS IP. **(C)** Origin firing efficiency for DONSON^AF647^ compared to mock-depleted and DONSON-depleted reactions (N – number of DNA molecules, error bars – 95% CI). Origins were counted using the Fen1^mKikGR^ channel. **(D)** Kymograms showing transient DONSON^AF647^ binding (green) during replication initiation. Dashed boxes mark productive binding, white arrows - non-productive binding. DONSON fluorescent labeling efficiency was ∼46% (Figure S4D). A biochemical replication assay was used to verify that recombinant DONSON efficiently supported DNA replication (Figure S4E-G). **(E)** Probability of productive (green) versus non-productive (gray) DONSON^AF647^ binding events (error bars – 95% CI). **(F)** Distributions of measured time delays between protein binding and origin firing for DONSON and GINS measured in separate experiments (blue bar marks the median, gray box – 95% CI for the median). P-values were computed using the two-sample Kolmogorov-Smirnov est. **(G)** Kymograms illustrating the relative timing of GINS^AF647^ and DONSON^AF546^ recruitment. At some origins, only one GINS molecule is fluorescently labeled. The disappearance of GINS^AF647^ is due to photobleaching. **(H)** Distribution of measured time delay between GINS recruitment and DONSON binding (time is shown in movie frames, 1 frame = 10 seconds; N – number of origins where both GINS and DONSON were detected; error bars – 95% CI estimated via bootstrapping).

To directly interrogate DONSON’s function, we visualized origin firing in a DONSON-depleted reaction supplemented with 10 nM of DONSON^AF647^ (Figure 4C, S4D-G). Like TopBP1 and RecQL4, DONSON^AF647^ transiently bound to DNA shortly before replication initiation (Figure 4D, green boxes, S4H), while a minority of DONSON binding events failed to trigger origin firing (Figure 4D, white arrows, Figure 4E). More importantly, the delay between DONSON binding and origin firing was nearly identical to that measured for GINS (Figure 4F), consistent with the idea that DONSON facilitates GINS recruitment.

If DONSON is the GINS dimerization scaffold, two copies of GINS and two copies of DONSON should be simultaneously recruited to the dormant origin. To test this hypothesis, we depleted GINS and DONSON from extract, supplemented the reaction with 10 nM DONSON^AF546^ and 30 nM GINS^AF647^, and simultaneously visualized their recruitment. Both GINS and DONSON signals were clearly detectable in 27 origin firing events (Figure 4G), and in ∼80% of them DONSON and GINS were recruited to DNA at the same time (Figure 4H). This observation supports the model where a DONSON dimer ensures the simultaneous delivery of two GINS molecules to the pre-RC (Figure 1G).

A recent study proposed that DONSON also mediates Cdc45 recruitment to origins ^17^. Unlike GINS, Cdc45 did not co-precipitate with DONSON (Figure 4A), suggesting that a different dimerization scaffold is responsible for recruiting two copies of Cdc45.

### Attempts to Recruit a Single Copy of GINS or Cdc45 Are Rapidly Rejected

Stable recruitment of two GINS^AF647^ molecules was often preceded by short-lived docking events (Figure 5A, red arrows, Figure 5B) that contained a single copy of GINS^AF647^ (Figure 5C) and lasted only ∼10 seconds on average (Figure 5D). This observation suggests that the initiation machinery can distinguish the recruitment of a single GINS (which binds weakly and quickly dissociates) from the simultaneous recruitment of two GINS – which bind tightly and are ultimately incorporated into a pair of CMG helicases (Figure 5E). It is unclear if these short-lived GINS docking events are mediated by a DONSON scaffold, or whether GINS alone can weakly bind the pre-RC. Similar short-lived docking was observed in the Cdc45 experiment (Figure 5F-H), suggesting that a similar mechanism rejects intermediates containing only one Cdc45 molecule (Figure S5).

**Figure 5:**
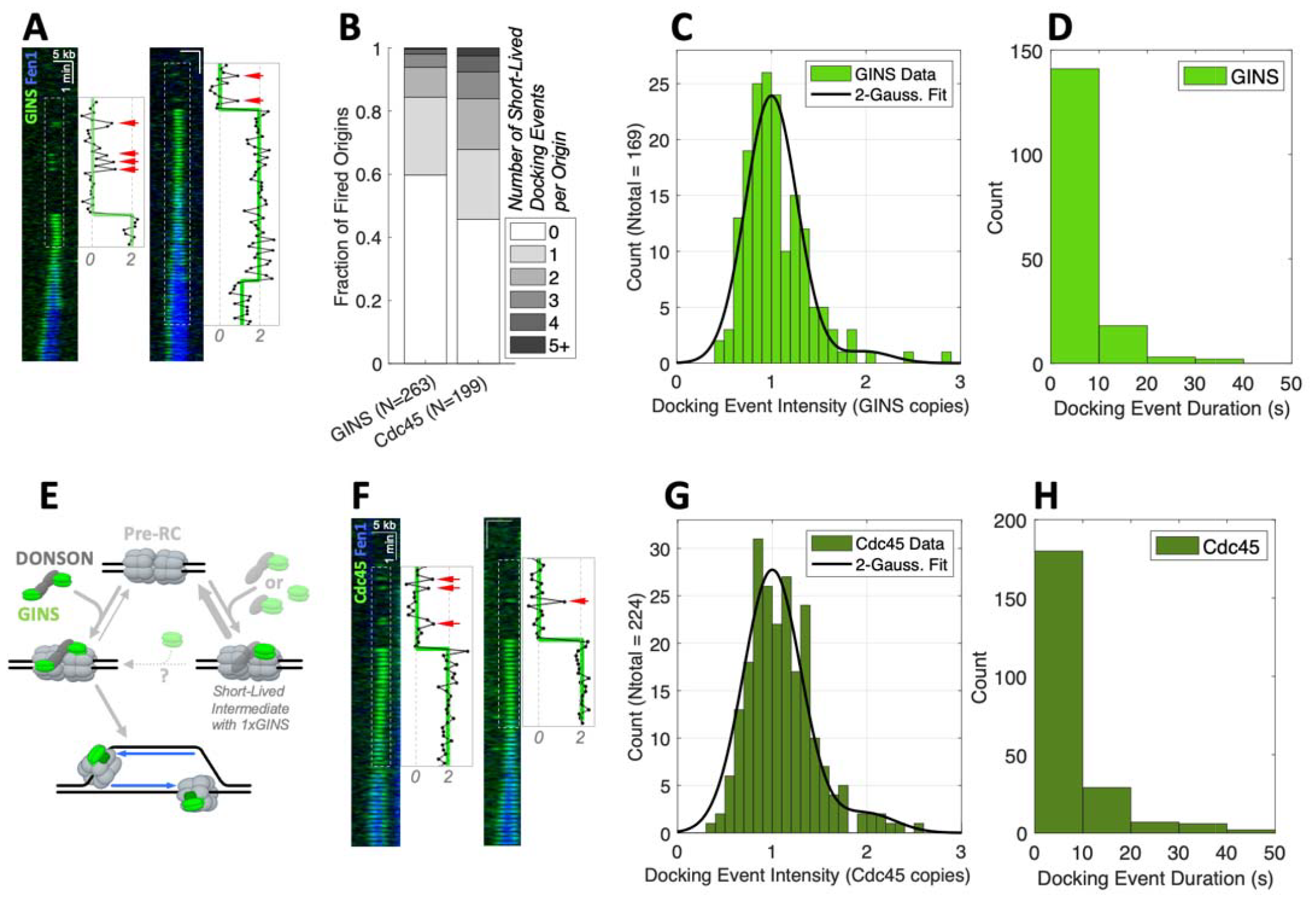
Attempts to Recruit a Single Copy of GINS or Cdc45 Are Rapidly Rejected. **(A)** Kymograms showing short-lived docking of GINS^AF647^ before origin firing (inset, red arrows). **(B)** The fraction of fired origins that were preceded by 0, 1, 2, 3, 4, or more short-lived GINS or Cdc45 docking events. Since the fluorescent labeling efficiency was ∼50% and ∼70% for GINS and Cdc45 respectively, w under-estimated the frequency of short-lived docking events. **(C)** Integrated intensity distribution for short-lived GINS docking events. The signal was normalized to the intensity of a single GINS molecule measured when the origin fired. The distribution was fit to the sum of two gaussians centered at 1.0 and 2.0 respectively. **(D)** The distribution of short-lived GINS docking event durations. **(E)** Kinetic model depicting the short-lived non-productive recruitment of a single GINS molecule (alone, with a DONSON monomer, or with a DONSON dimer) versus the productive recruitment of two GINS copies facilitated by a DONSON dimer. Thicker arrows depict faster rate constants. **(F)** Kymograms illustrating short-lived docking of Cdc45^AF647^ before origin firing (inset, red arrows). **(G)** Integrated intensity distribution for short-lived Cdc45 docking events and the corresponding double Gaussian fit. **(H)** The distribution of short-lived Cdc45 docking event durations.

### RecQL4 Facilitates DONSON Dissociation and Promotes the Activation of Fully Assembled CMGs

In experiments where GINS^AF647^ recruitment was monitored, two molecules of GINS were often stably recruited to DNA but did not result in origin firing (Figure 6A, S6A). Such long-lived dual-GINS recruitment events were not observed when Cdc45 was depleted (Figure S6B). Similarly, two Cdc45 molecules were often stably recruited to DNA without triggering origin firing (Figure 6B, S6A), but not when GINS was depleted (Figure S6C). We hypothesize that non-productive dual-GINS or dual-Cdc45 binding states represent fully assembled CMG•CMG complexes that remain inactive.

**Figure 6:**
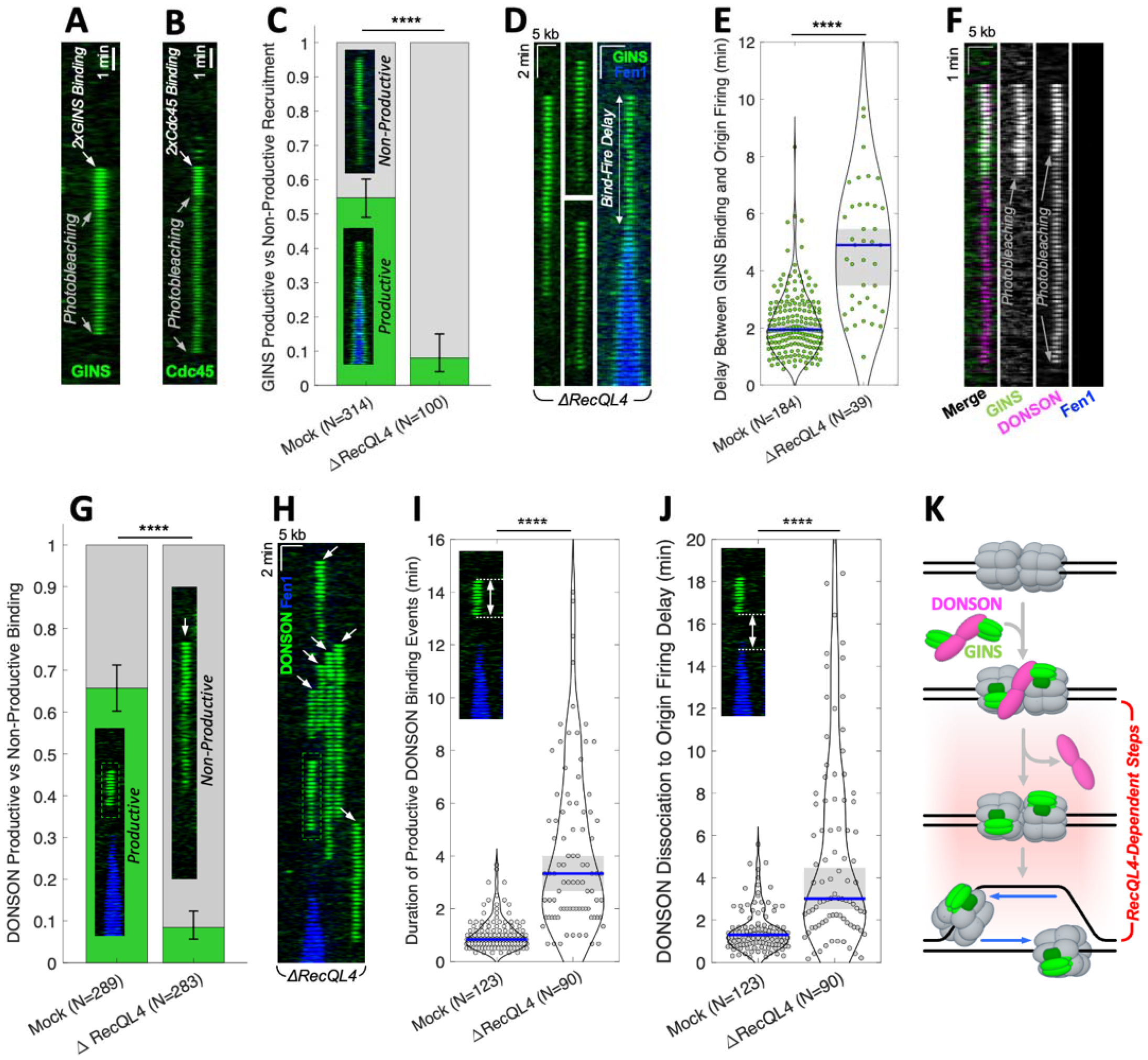
RecQL4 Facilitates DONSON Dissociation, Promotes the Activation of Fully Assembled CMGs. **(A)** Kymogram showing non-productive dual-GINS^AF647^ recruitment. **(B)** Kymogram showing non-productive dual-Cdc45^AF647^ recruitment. In most non-productive recruitment events, the two GINS/Cdc45 molecules are simultaneously recruited (Figure S6A). **(C)** Fraction of productive (green) and non-productive (gray) dual-GINS recruitment events in mock-depleted versus RecQL4-depleted reactions. **(D)** Kymograms showing non-productive and productive dual-GINS recruitment events in a RecQL4-depleted replication reaction. **(E)** Distributions of time delays between GINS recruitment and origin firing in mock-depleted versus RecQL4-depleted reactions. **(F)** Kymogram showing a typical non-productive dual-DONSON^AF546^ binding accompanied GINS^AF647^ recruitment. **(G)** Fraction of productive (green) and non-productive (gray) DONSON binding events in mock-depleted versus RecQL4-depleted reactions. Non-productive DONSON recruitment events a e much longer than productive ones (Figure S6D). **(H)** Kymogram illustrating DONSON binding to licensed DNA in RecQL4-depleted extract. Boxes and arrows indicate productive and non-productive DONSON binding. Occasionally, initiation intermediates containing stably bound DONSON are pushed by progressing forks (Figure S6E). **(I)** Duration of productive DONSON binding events in mock-depleted versus RecQL4-depleted reactions. **(J)** The time delay between DONSON dissociation and origin firing in mock-depleted versus RecQL4-depleted reactions. **(K)** Cartoon illustrating that DONSON release after successful GINS recruitment and CMG helicase activation are RecQL4-dependent steps during initiation. In panels **(C)** and **(G)** error bars indicate 95% CI. In panels **(E), (I)**, and **(J)**, blue bars indicate median values, gray boxes show the 95% CI for the median. To minimize photobleaching, experiments in panels **(D)** and **(H)** were conducted at 20 seconds/frame resolution. P-values were computed using the two-sample Kolmogorov-Smirnov test.

RecQL4 transiently bound dormant origins after GINS recruitment (Figure 3E), suggesting that RecQL4 plays a role in CMG activation. Some replication initiation occurred in RecQL4-depleted extract (Figure 3B) even though the depletion was very efficient (Figure S3E). Origin firing was presumably driven by trace amounts of RecQL4 that evaded depletion. This enabled us to measure the properties of productive and non-productive DONSON binding events when RecQL4 activity was rate-limiting and origin firing was very inefficient.

Depleting RecQL4 led to a dramatic increase in non-productive dual-GINS recruitment events (Figure 6C-D). More importantly, when origins did fire, the delay between GINS binding and origin firing was ∼2.5-fold longer in RecQL4-depleted extract (Figure 6E), consistent with the hypothesis that RecQL4 facilitates CMG activation.

In experiments where both GINS^AF647^ and DONSON^AF546^ were visualized, long-lived non-productive GINS recruitment was accompanied by long-lived binding of two DONSON molecules (Figure 6F). This observation suggests that after delivering GINS to the pre-RC, DONSON remains stably bound to inactive helicases. Therefore, DONSON must dissociate from the dormant origin before the two CMG helicases can be activated and replication can initiate. Consistent with this idea, depleting RecQL4 led to a large increase in non-productive DONSON binding events (Figure 6G-H).

Depleting RecQL4 also led to a large increase (∼3-fold) in the median duration of productive DONSON binding events (Figure 6I), suggesting that RecQL4 facilitates DONSON dissociation from assembled helicases. Finally, depleting RecQL4 led to a significant increase (∼2-fold) in the time delay between DONSON dissociation and origin firing (Figure 6J), suggesting that RecQL4 also promotes CMG helicase activation following DONSON release (Figure 6K).

### A Monomer of TopBP1 is Sufficient to Promote Origin Firing

The symmetry required for bidirectional replication is enforced at several stages: pre-RCs contain two Mcm2-7 complexes^1,2^, two GINS molecules are recruited at once, and two copies of Cdc45 are recruited simultaneously. We asked if TopBP1 helps enforce the two-fold symmetry of replication initiation intermediates.

The photobleaching assay revealed that the vast majority of full-length TopBP1 and truncated TopBP1 are monomeric *in vitro* (Figure S7A). However, TopBP1 may be able to oligomerize in nuclear extract, in the presence of other proteins. To determine TopBP1 stoichiometry during initiation, we measured the intensity of productive binding events in the trunc-TopBP1^AF647^ experiment. A significant fraction (∼40%) of origin firing events was facilitated by a single copy of trunc-TopBP1^AF647^ (Figure 7A). When two trunc-TopBP1^AF647^ copies were detected, their recruitment was sequential in most cases (Figure 7B), suggesting that trunc-TopBP1 did not form stable dimers in extract. These results indicate that a single copy of trunc-TopBP1 is sufficient to support bidirectional origin firing and suggest that TopBP1 does not enforce two-fold symmetry during initiation.

**Figure 7:**
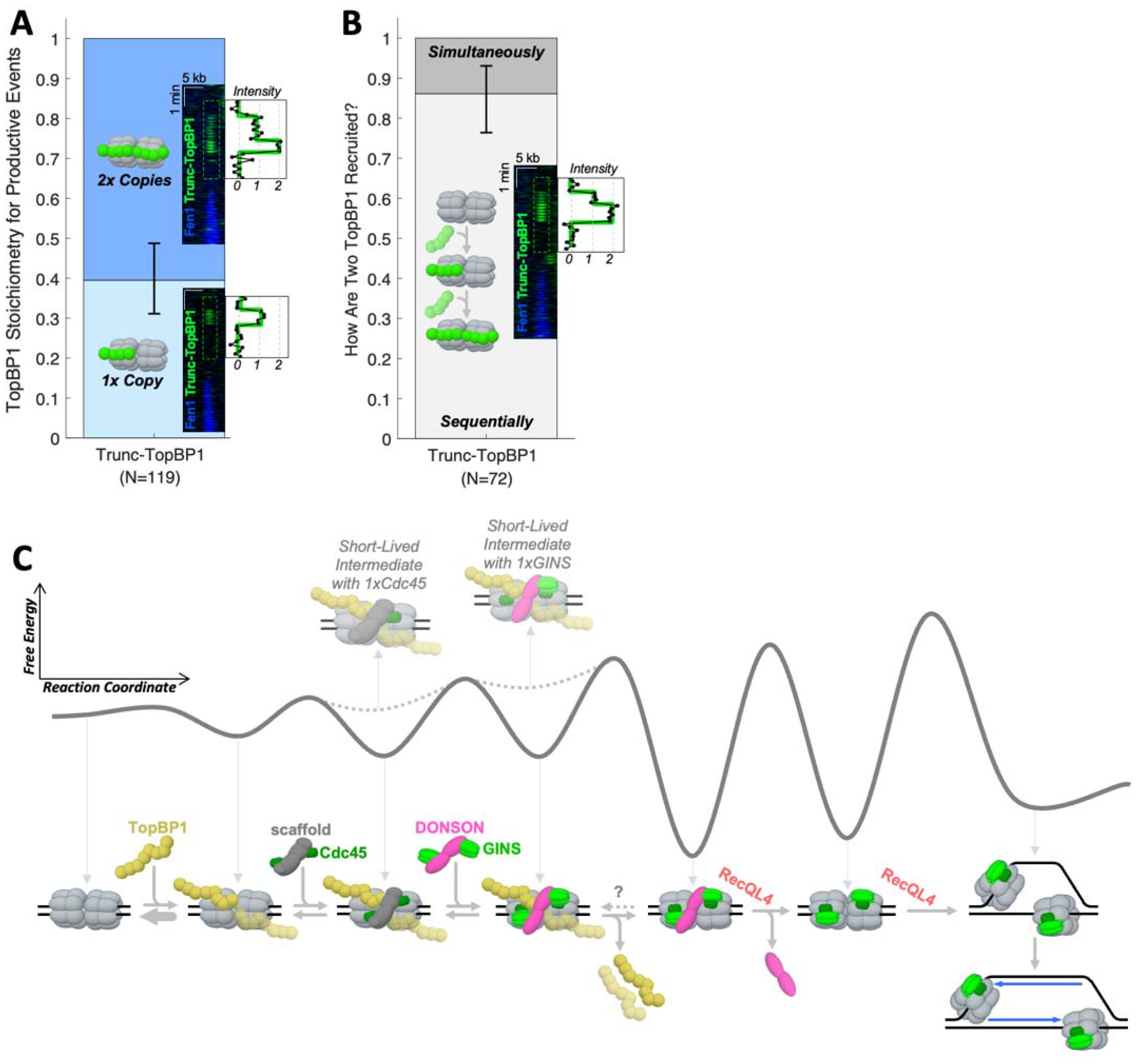
Model of Bidirectional Replication Initiation in Metazoa. **(A)** Fraction of productive binding events that contain one or two copies of trunc-TopBP1^AF647^ (error bars - 95% CIs). Insets show example kymograms with integrated signal intensity. **(B)** Of all productive binding events with two copies of trunc-TopBP1^AF647^, how many are recruited sequentially versus simultaneously? (error bars - 95% CIs). **(C)** Model of metazoan replication initiation based on single-molecule imaging data presented in this study. The thick gray curve is a qualitative depiction of the energy landscape for the proposed reaction scheme, dashed curves illustrate the energy level corresponding to short-lived intermediates with a single copy of Cdc45 or GINS. Treslin-MTBP, Mcm10, and other replication proteins are not shown for clarity. The timing of dissociation for the Cdc45 dimerization scaffold is not known and not shown. DONSON dissociation and CMG activation depend on RecQL4 and are most likely irreversible.

## DISCUSSION

### A New Model of Metazoan Replication Initiation

In this study we leveraged single-molecule imaging to visualize replication initiation in real time. Our experiments revealed a surprisingly dynamic process governed by a network of transient protein-protein interactions which previously confounded the interpretation of ensemble experiments. Our observations suggest a new model of replication initiation where several short-lived intermediates are needed to correctly assemble and activate two CMG helicases at each origin (Figure 7C).

Proper assembly of two replicative helicases at each origin is achieved via the simultaneous recruitment of two Cdc45 molecules, and briefly afterward – of two GINS molecules. The delivery of these proteins to the origin and their incorporation into replicative helicases is carefully orchestrated by several firing factors. By directly visualizing three key firing factors (TopBP1, DONSON, and RecQL4), we revealed that all three firing factors transiently dock to the dormant origin but do not travel with the replication fork. This paints a clear picture of their function and reconciles previous conflicting reports.

Single-molecule imaging suggests that TopBP1 is recruited to Mcm2-7 before Cdc45 or GINS. Consistent with this conclusion, careful quantification of GINS co-immunoprecipitation from egg extract indicates that TopBP1 does not form a stable stoichiometric complex with DONSON and GINS (Figure 4B). These findings are inconsistent with the currently accepted pre-LC model of initiation ^16,17,21^. Once bound to the dimer of Mcm2-7, TopBP1 acts as a “docking adapter” to recruit the GINS•DONSON•DONSON•GINS complex from solution. Since TopBP1 contains several BRCT phospho-peptide binding modules, we hypothesize that TopBP1 acts as “reader” of Mcm2-7 phosphorylation. For example, TopBP1 may bind phosphorylated pre-RCs more stably, increasing the likelihood of origin firing. Data presented here indicate that TopBP1 dimerization is not required for replication initiation (Figure 7A). It is possible that TopBP1 dimerizes only at high concentrations but remains monomeric at low concentrations, which are nevertheless sufficient to support origin firing. Consistent with this idea, knocking down TopBP1 in human cells did not impair DNA replication, but fully supressed ATR signalling ^9^ which requires TopBP1 oligomers ^7,8,31^.

DONSON acts as the dimerization scaffold for GINS, ensuring the simultaneous delivery of two copies of GINS to the Mcm2-7 double hexamer. DONSON was initially thought to travel with replication forks and stabilize them in response to DNA damage ^18,19^. Single-molecule imaging revealed that DONSON is not incorporated into active replisomes. DONSON may protect genome integrity by promoting dormant origin firing near stalled replication forks. We surmise that biochemical assays used in previous studies detected the long-lived CMG•DONSON•DONSON•CMG intermediate. It is unclear how this inactive complex is removed from chromatin – via passive dissociation or active extraction (perhaps through a ubiquitin-dependent pathway). How the helicase is activated remains one of the most mysterious aspects of DNA replication. Single-molecule imaging indicates that RecQL4 plays a key role in this process: it promotes DONSON release from fully assembled CMGs and facilitates helicase activation after DONSON dissociation. How RecQL4 accomplishes this is unknown: it may unwind dsDNA at the origin, it may remodel the head-to-head CMG dimer, or it may recruit Mcm10 as proposed previously ^32^.

The model of initiation outlined in Figure 7C appears to be a classic example of Hopfield kinetic proofreading^33–35^ – wherein a multi-step biochemical reaction has one or more reversible steps and at least one irreversible step leading to the correct final product. Incorrect intermediate products prematurely exit this pathway, whereas correct intermediates successfully complete the reaction. Kinetic proofreading ensures high fidelity in multi-step biochemical reactions like tRNA aminoacylation^36^, T-cell receptor signalling^37^, homologous recombination^38^, and SOS response activation in bacteria^39^.

In the replication initiation reaction scheme, we identified at least three reversible steps: TopBP1 binding, Cdc45 recruitment, and GINS recruitment (Figure 7C). These reversible steps discard incomplete or defective protein assemblies. For example, attempts to recruit a single copy of GINS or Cdc45 are very short lived and do not promote origin firing. We also identified two steps that are most likely irreversible: DONSON eviction from the CMG•CMG dimer and CMG helicase activation, both of which are RecQL4-dependent (Figure 7C). These irreversible steps commit the dormant origin to firing. More importantly, these irreversible steps are critical to ensuring the high fidelity required to initiate DNA replication bi-directionally from tens of thousands of origins during each cell division.

### Differences Between Metazoa and Budding Yeast

Origin firing was previously reconstituted with purified yeast proteins ^40^, but not with human proteins ^41^, suggesting the existence of additional regulators of initiation in metazoa. One of them was recently identified in the form of DONSON. Using InterPro, thousands of putative DONSON homologs can be found in metazoa, plants, algae, amoeba, and basal fungi, but not in dikarya - the taxa containing budding yeast ^42^. This suggests that DONSON was present in the last eukaryotic common ancestor but was lost in *S. cerevisiae*.

Our observation that two Cdc45 molecules simultaneously bind to the pre-RC was surprising because in budding yeast two Cdc45 molecules are sequentially recruited to Mcm2-7 ^43^. This likely reflects real differences in how initiation is regulated in yeast versus metazoa. Since *S. cerevisiae* has ∼500 origins versus 10^4^-10^5^ in human cells, it is possible that yeast achieves robust bi-directional replication with fewer layers of symmetry-enforcing regulation.

The Cdc45 dimerization scaffold postulated here remains to be identified. It may be another metazoan-specific initiation factor. Alternatively, Cdc45 dimerization could be mediated by Treslin-MTBP, which can oligomerize ^44,45^ and can interact with Cdc45 ^46^.

### Limitations of The Study

Xenopus egg extracts contain very high concentrations of replication factors, which are needed to support several fast cell divisions following egg fertilization ^47^. Mock-depleted replication reactions contained ∼200 nM of GINS, ∼200 nM of Cdc45, ∼150 nM of TopBP1, ∼250 nM of RecQL4, and ∼300 nM of DONSON. To maintain an acceptable signal-to-noise ratio, the concentration of fluorescent proteins was capped at 30 nM. Our experiments show that the mechanism of replication initiation is robust across a wide range of GINS, Cdc45, TopBP1, RecQL4, and DONSON concentrations. It is unclear how concentrations used in this study compare to physiological conditions, as there are no accurate measurements of replication protein concentrations in cells. Estimates from the Protein Abundance Database (pax-db.org) vary widely among cell types: 50-500 nM for GINS, 20-100 nM for Cdc45, 5-50 nM for TopBP1, 10-50 nM for DONSON, and 1-10 nM for RecQL4.

To track CMG movement after origin firing and to resolve protein binding to different locations on DNA, the DNA substrate was stretched to ∼80% of its contour length, therefore it could not be fully chromatinized. We concluded that TopBP1, DONSON, and RecQL4 do not undergo long-range diffusion along DNA during initiation. However, given the spatial resolution of our assay (∼300-500 bp), we cannot rule out that some proteins (for example RecQL4, which has a helicase domain) may travel short distances near the dormant origin.

To date we have not found an oxygen scavenging system that does not inhibit replication in extract. To minimize photobleaching, in most experiment the image acquisition rate was limited to 10 s/frame, resulting in a half-life of 4-5 min for Alexa Fluor 647 (Figure S7B). Photobleaching limited our ability to accurately measure the duration of long-lived events. However, the effect of photobleaching was minimal on productive binding events for TopBP1, DONSON, and RecQL4, which lasted for only 1-2 min.

We spent considerable effort optimizing fluorescent labeling of recombinant proteins to maximize labeling efficiency and minimize any effect of labeling on protein function. TopBP1 and RecQL4 were very efficiently labeled (>90%), whereas GINS, and Cdc45 were labeled less efficiently (∼50% and ∼70% respectively). As a result, we focused our analysis on initiation events where both copies of GINS or Cdc45 were labeled and visible. Due to incomplete labeling (∼46%), some DONSON binding events were invisible, while others contained only one labeled DONSON copy.

To establish the recruitment order of GINS, and Cdc45, we separately measured the time delay between protein recruitment and origin firing. These measurements were remarkably consistent between different extract preparations (Figure S7C-D) and between experiments with the same extract (Figure S7E). We relied on similar time delay measurements to infer that TopBP1 was recruited before Cdc45 and GINS. Although we could only interpret a fraction of full-length TopBP1^AF647^ binding events (Figure S3C), the full length and truncated TopBP1 constructs exhibited nearly identical binding-dissociation kinetics (Figure S7F), suggesting that the truncated construct is an excellent surrogate for the full-length protein.

## Acknowledgments

We thank members of the Chistol, Cimprich, and Ferrell labs for suggestions and feedback; Karlene Cimprich, James Ferrell, and Agnel Sfeir for critical reading of the manuscript; William Dunphy and Akiko Kumagai for *Xenopus* topbp1 cDNA.; and Johannes Walter for sharing unpublished results. R.T. was supported in part by the Stanford School of Medicine Dean’s fellowship. S.E.B. is supported by the Stanford Graduate Fellowship and the New Science fellowship. L.A.S. is supported in part by a Blavatnik Family Fellowship Fund. G.C. is supported by an NSF CAREER Award (2144481), an NIGMS R35 award (GM147060), and an American Cancer Society seed grant (228425).

## Author Contributions

G.C., R.T., and S.E.B. wrote the manuscript with input from all authors. G.C. conceived and directed the project. G.C., R.T., and S.E.B. conceptualized and designed most experiments. R.T. prepared all recombinant proteins. R.T. and S.E.B. performed biochemical assays in *X*.*l*. extract and single-molecule experiments. R.T. and L.A.S. performed immunoprecipitation assays. S.E.B. raised and validated the Cdc45 custom antibody. D.S. raised and validated TopBP1 custom antibodies. R.T. raised and validated DONSON and RPA custom antibodies. L.A.S. raised and validated RecQL4 custom antibodies. G.C. wrote MATLAB code to analyze single-molecule experiments. G.C., R.T. and S.E.B. analyzed single-molecule data.

