## Supplementary Figures for "Single-Molecule Imaging Reveals the Mechanism of Bidirectional Replication Initiation in Metazoa"

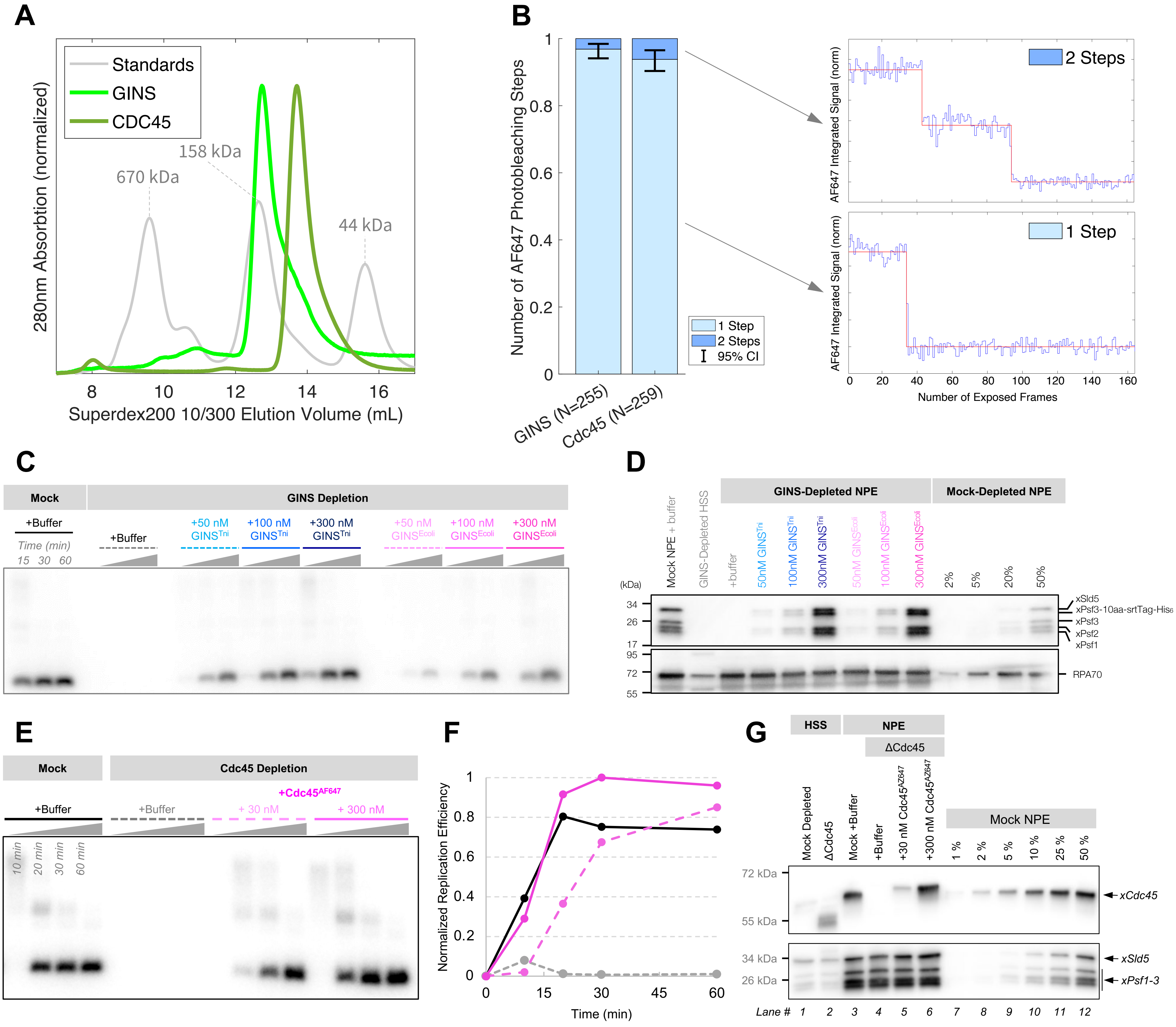


**Figure S1: Validation and Characterization of Recombinant GINS and Cdc45, related to Figure 1.**
**(A)** Gel filtration chromatogram of recombinant GINS, Cdc45, and gel filtration standards. **(B)** The oligomeric state of GINS and Cdc45 was measured using an in vitro photobleaching assay. Left: bar plots showing the probability that spots corresponding to labeled GINS/Cdc45 photobleached in one step (light blue) or two steps (dark blue). Right: representative traces illustrating 1-step and 2-step photobleaching. N – number of fluorescent spots analyzed, error bars – 95% CI estimated via bootstrapping. **(C)** Plasmid replication assay comparing the biochemical activity of mock-depleted extract versus GINS-depleted extract supplemented with buffer, 50-300 nM of recombinant GINS purified from insect cells (Tni) or bacteria (*E. coli)*. GINS purified from bacteria was roughly half as active as GINS purified from insect cells. **(D)** Immunoblots of samples corresponding to reactions from panel **(C)**. RPA70 was used as a loading control. The replication reaction consists of equal parts HSS, NPE, and buffer. Based on this blot, the estimated concentration of GINS is ~200 nM in the replication reaction and ~600 nM in undiluted NPE. **(E)** Plasmid replication assay comparing the biochemical activity of mock-depleted extract versus GINS-depleted extract supplemented with buffer or with 30-300 nM of fluorescently labeled recombinant $Cdc45^{AF647}$. **(F)** Plot of integrated signal intensities from panel **(E)** with matching line styles and colors. **(G)** Immunoblots of samples corresponding to reactions from panel **(E)**. GINS was used as a loading control.

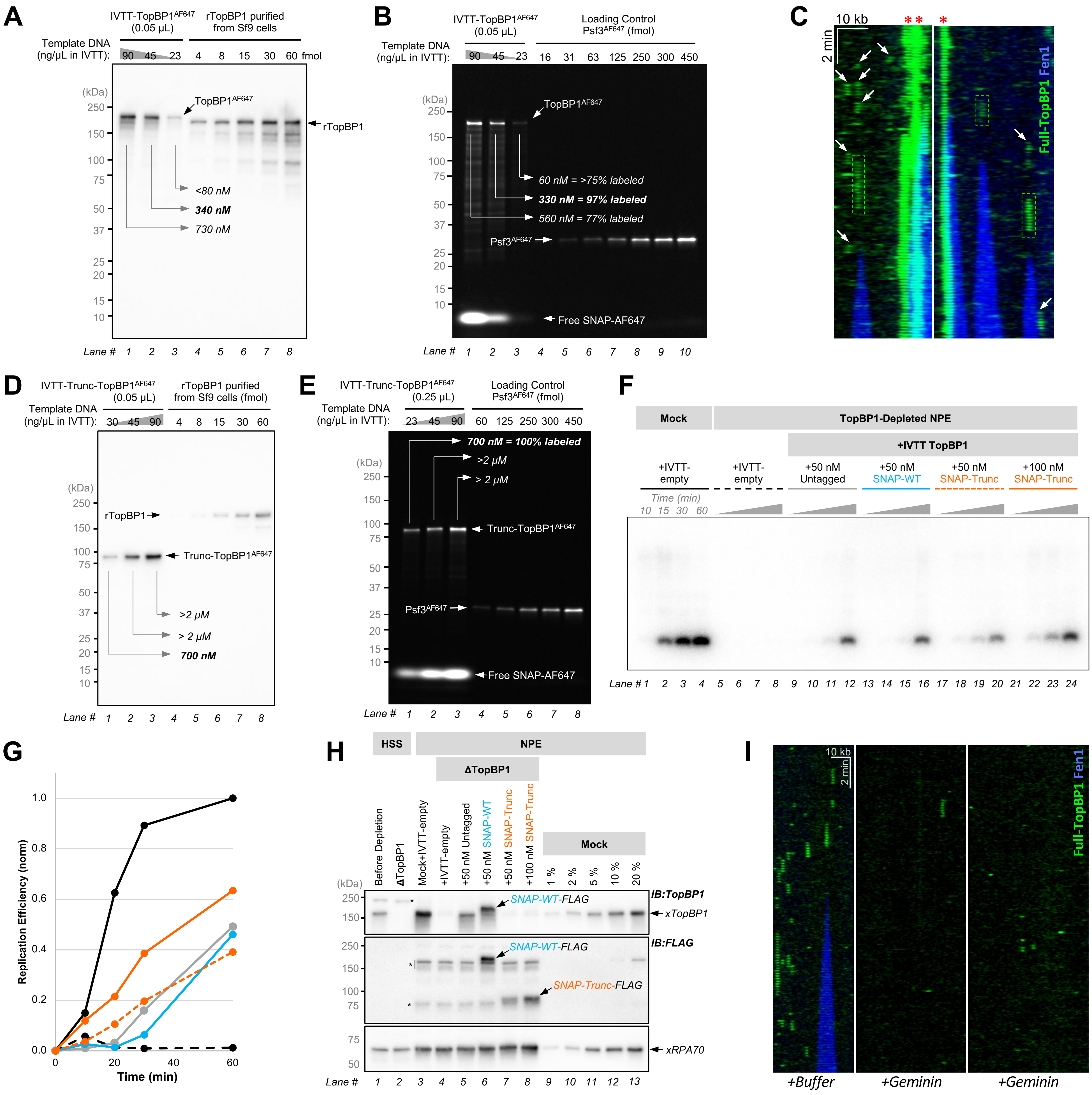


**Figure S2: Validation and Characterization of Recombinant TopBP1, related to Figure 2**

**(A)** SNAP-tagged TopBP1 was expressed via In Vitro Transcription-Translation (IVTT) by using the indicated concentration of template plasmid in IVTT reactions. SNAP-tagged TopBP1 expressed in IVTT was labeled with Alexa Fluor 647 (AF647) by incubating the IVTT reaction with SNAP-Surface Alexa Fluor 647 at 4 °C overnight. IVTT-TopBP1$^{AF647}$ was separated by SDS-PAGE, and TopBP1 concentration was estimated via semi-quantitative western blotting. **(B)** IVTT-TopBP1$^{AF647}$ fluorescent labeling efficiency was estimated by comparing the AF647 signal with known amounts of GINS$^{AF647}$ where the fluorophore is on Psf3. The 330 nM TopBP1$^{AF647}$ preparation (lane 2) was used for single-molecule assays. **(C)** Representative kymograms from a single-molecule experiment with 10 nM of full-length TopBP1$^{AF647}$. Boxes indicate productive TopBP1 binding events, arrows show non-productive events, red asterisks mark large TopBP1 assemblies that may represent liquid-liquid phase separation (LLPS). **(D)** Semi-quantitative immunoblot of truncated TopBP1 expressed via IVTT. **(E)** Estimation of trunc-TopBP1 fluorescent labeling efficiency. The 700 nM Trunc-TopBP1$^{AF647}$ preparation (lane 1) was used for single-molecule assays. **(F)** Plasmid replication assay comparing the biochemical activity of mock-depleted reaction (lanes 1-4) versus TopBP1-depleted reaction supplemented with buffer (lanes 5-8), or TopBP1 constructs expressed in IVTT: 50 nM of untagged TopBP1 (lanes 9-12), 50 nM of SNAP-tagged full-length wildtype TopBP1 (lanes 13-16), and 50 nM or 100 nM of SNAP-tagged truncated TopBP1 (lanes 17-20 and lanes 21-24 respectively). **(G)** Plot of integrated signal intensities from panel **(F)**. 50 nM (~20% of endogenous TopBP1 concentration) of untagged (solid gray), SNAP-tagged full length (solid blue), and SNAP-tagged truncated TopBP1 (dashed orange) rescue DNA replication to ~50% level of the mock reaction. **(H)** Immunoblots of samples corresponding to reactions from panel **(F)**. **(I)** Licensing DNA in HSS supplemented with 300 nM geminin (which inhibits pre-RC loading) greatly reduced trunc-TopBP1 binding in the single-molecule assay.

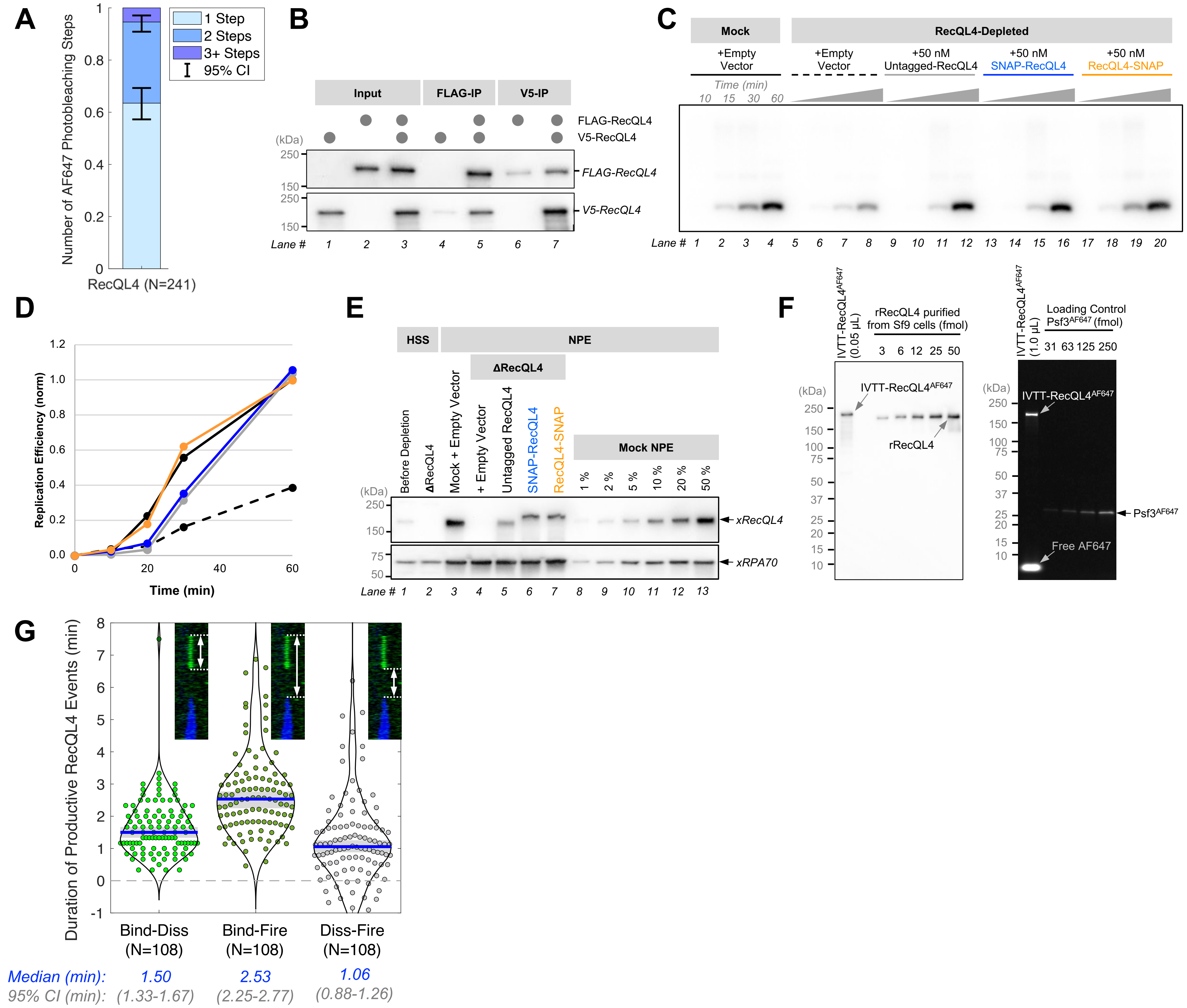


**Figure S3: Validation and Characterization of Recombinant RecQL4, related to Figure 3**

**(A)** The oligomeric state of IVTT-RecQL4$^{AF647}$ was measured using an in vitro photobleaching assay. Bar plot depicts the probability that spots corresponding to labeled RecQL4 photobleached in one step (light blue), two steps (medium blue), or more steps (dark blue). N – number of fluorescent spots analyzed, error bars – 95% CI estimated via bootstrapping. **(B)** Co-IP experiment of FLAG-RecQL4 and V5-RecQL4. N-terminally V5-tagged RecQL4 (lanes 1, 3 – 5, 7) and N-terminally FLAG-tagged RecQL4 (lanes 2, 3, 5 – 7) were expressed via IVTT, and immunoprecipitated using anti-FLAG beads (lanes 4 and 5) or anti-V5 beads (lanes 6 and 7). Samples were separated by SDS-PAGE, transferred to a PVDF membrane, and blotted with anti-FLAG (upper panel) or anti-V5 antibody (lower panel). V5-RecQL4 co-precipitated with FLAG-RecQL4 (lanes 4 and 5), and vice versa (lanes 6 and 7), suggesting that RecQL4 forms oligomers. **(C)** Plasmid replication assay comparing the biochemical activity of mock-depleted extract (lanes 1-4) versus RecQL4-depleted extract supplemented with buffer (lanes 5-8), 50 nM of untagged RecQL4 (lanes 9-12), 50 nM of N-term SNAP-tagged RecQL4 (lanes 13-16), or 50nM of C-term SNAP-tagged RecQL4 (lanes 17-20). **(D)** Plot of integrated signal intensities from panel **(C)** with matching line styles and colors. **(E)** Immunoblots of samples corresponding to reactions from panel **(C)**. 50 nM RecQL4 corresponds to ~15% of endogenous RecQL4 concentration. RPA70 was used as a loading control. **(F)** Estimation of RecQL4$^{AF647}$ fluorescent labeling efficiency. C-terminally SNAP tagged RecQL4 was expressed via IVTT and incubated with SNAP-Surface Alexa Fluor 647 at 4 °C overnight. Left: RecQL4 concentration was estimated by semi-quantitative western blotting. Right: RecQL4 fluorescent labeling efficiency was estimated by comparing the AF647 signal with known amounts of GINS$^{AF647}$ where the fluorophore is on Psf3. This preparation yielded ~210 nM of RecQL4, of which ~200 nM was fluorescently labeled (~95% labeling efficiency). **(G)** The distribution of measured time delays for productive RecQL4 binding events: RecQL4 binding to RecQL4 dissociation delay (left), RecQL4 binding to origin firing delay (middle), and RecQL4 dissociation to origin firing delay (right). Blue lines – median values, gray boxes – 95% CIs estimated via bootstrapping.

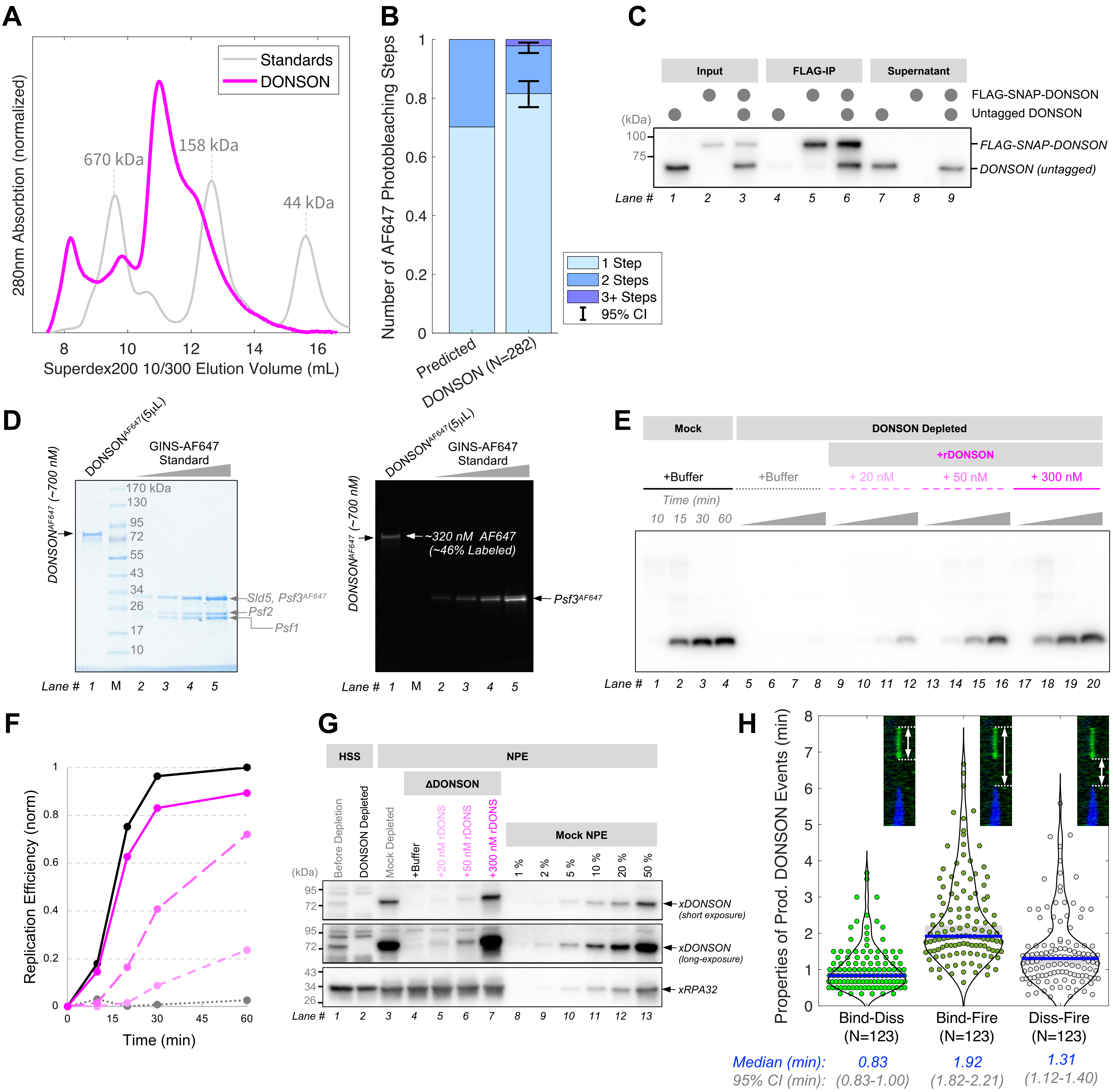


**Figure S4: Validation and Characterization of Recombinant DONSON Purified from Sf9 cells, related to Figure 4**

**(A)** Gel filtration chromatogram of recombinant DONSON with standards. **(B)** In vitro photobleaching analysis of recombinant DONSON, of which ~46% was labeled with Alexa Fluor 647. Left bar: the predicted probability of 1-step and 2-step photobleaching events given the ~46% labeling efficiency under the assumption that DONSON forms stable dimers in vitro. Right bar: the measured probability that DONSON$^{AF647}$ spots photobleached in one, two, or more steps. N – number of fluorescent spots analyzed, error bars – 95% CI estimated via bootstrapping. **(C)** Co-IP experiment of untagged DONSON and N-terminally FLAG-SNAP-tagged DONSON (DONSON-SNAP-FLAG). Untagged DONSON (lanes 1, 3, 4, 6, 7, and 9) and FLAG-SNAP-DONSON (lanes 2, 3, 5, 6, 8, and 9) were expressed by IVTT, and were immunoprecipitated using anti-FLAG beads (lanes 4 – 6). Samples were separated by SDS-PAGE, transferred to a PVDF membrane, and blotted with anti-DONSON antibody. Untagged DONSON co-precipitated with FLAG-SNAP-DONSON, suggesting that DONSON forms oligomers. **(D)** Left: DONSON$^{AF647}$ was separated by SDS-PAGE and stained with Coomassie brilliant blue. Right: DONSON fluorescent labeling efficiency was estimated by comparing the AF647 signal with known amounts of GINS$^{AF647}$ where the fluorophore is on Psf3. This preparation yielded ~700 nM of DONSON, of which ~320 nM was fluorescently labeled (~46% labeling efficiency). **(E)** Plasmid replication assay comparing the biochemical activity of mock-depleted extract (lanes 1-4) versus DONSON-depleted extract supplemented with buffer (lanes 5-8), 20 nM (lanes 9-12), 50 nM (lanes 13-16), or 300 nM (lanes 17-20) of recombinant DONSON. **(F)** Plot of integrated signal intensities from panel **(E)** with matching line styles and colors. **(G)** Immunoblots of samples corresponding to reactions from panel **(E)**. 50 nM DONSON corresponds to ~5% of endogenous DONSON concentration in NPE. RPA32 was used as a loading control. **(H)** The distribution of measured time delays for productive DONSON binding events: DONSON binding to DONSON dissociation delay (left), DONSON binding to origin firing delay (middle), and DONSON dissociation to origin firing delay (right). Blue lines – median values, gray boxes – 95% CIs estimated via bootstrapping.

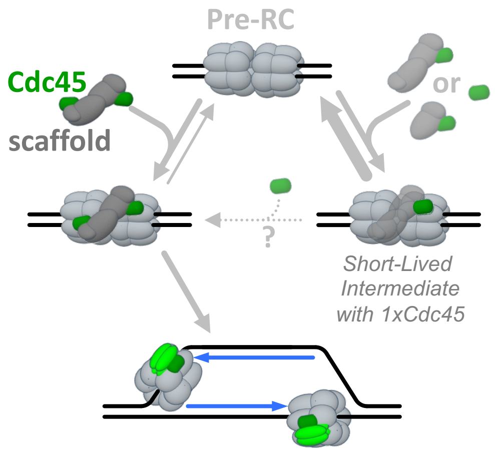


**Figure S5: Short-Lived Cdc45 Docking Events, related to Figure 5**
Kinetic model depicting the short-lived non-productive recruitment of a single Cdc45 molecule (alone, with a scaffold monomer, or with a scaffold dimer) versus the productive recruitment of two Cdc45 copies facilitated by a scaffold dimer. Thicker arrows depict faster rate constants.

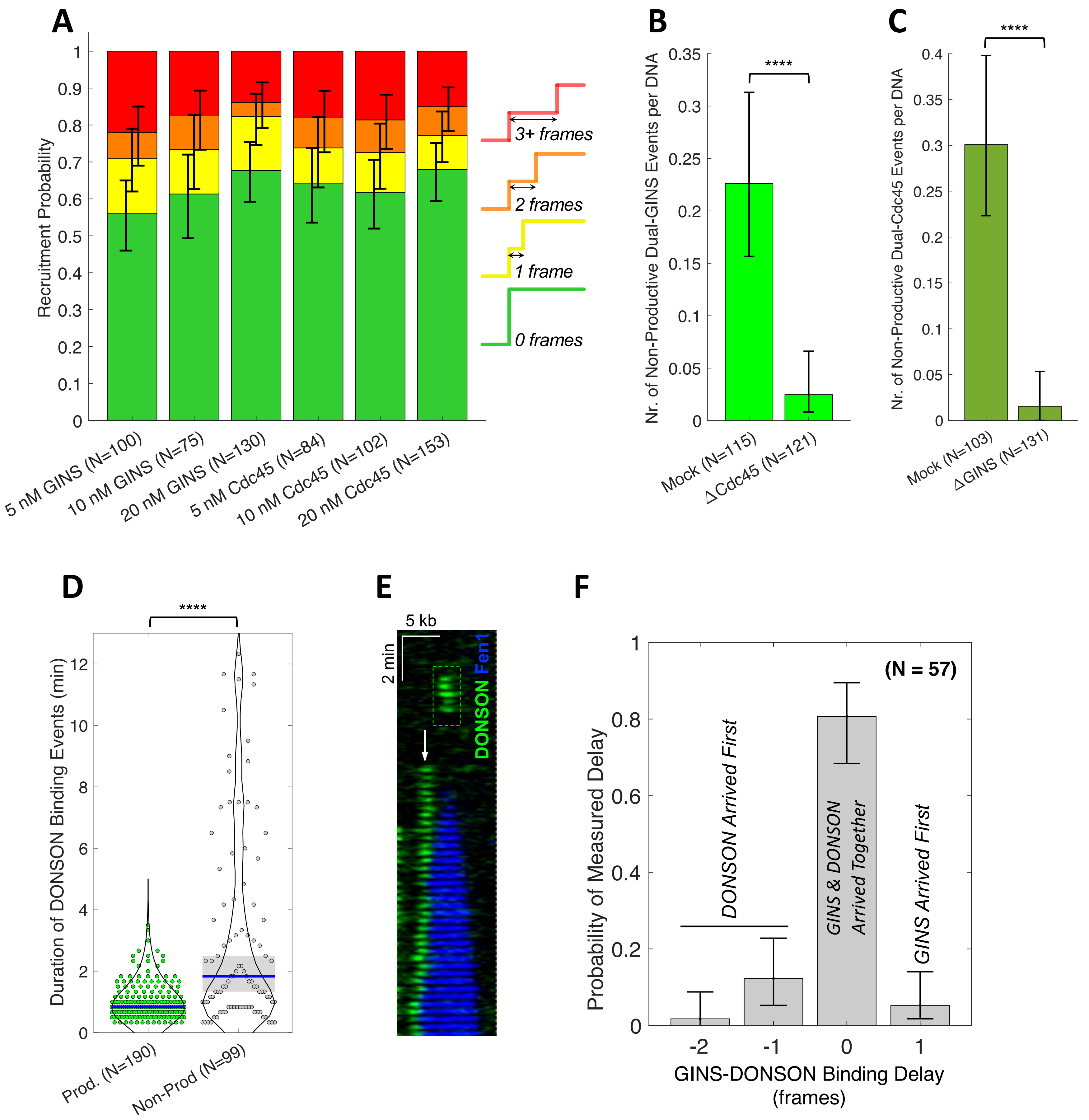


**Figure S6: Detailed Analysis of Long-Lived Non-Productive GINS, Cdc45, and DONSON Recruitment Events, related to Figure 6**
**(A)** Breakdown of how two copies of GINS and Cdc45 were recruited to long-lived non-productive events. In most events, two molecules of GINS/Cdc45 appeared simultaneously (0 frames, green). In the other events, two copies of $GINS^{AF647}$/$Cdc45^{AF647}$ showed up sequentially, where the second molecule appeared 1 frame (yellow), 2 frames (orange), or 3+ frames (red) after the first. N – number of long-lived non-productive binding events containing two copies of $GINS^{AF647}$ or $Cdc45^{AF647}$, error bars – 95% CIs estimated via bootstrapping. **(B)** The number of long-lived non-productive events containing two copies of $GINS^{AF647}$ observed per DNA molecule in mock-depleted (left) or Cdc45-depleted extract (right). N – number of DNA molecules analyzed, error-bars – 95% CIs estimated via bootstrapping. **(C)** The number of long-lived non-productive events containing two copies of $Cdc45^{AF647}$ observed per DNA molecule in mock-depleted (left) or GINS-depleted extract (right). N – number of DNA molecules analyzed, error-bars – 95% CIs estimated via bootstrapping. **(D)** The duration of productive and non-productive DONSON binding events. Blue bars and gray boxes represent medians and 95% CIs respectively. **(E)** Representative kymogram depicting a non-productive long-lived DONSON binding event (white arrow) being pushed by an active replication fork. **(F)** Distribution of measured time delays between GINS recruitment and DONSON binding for non-productive events. Time is shown in movie frames (1 frame = 10 seconds), N – number of non-productive events where both GINS and DONSON were detected, error bars – 95% CI estimated via bootstrapping.

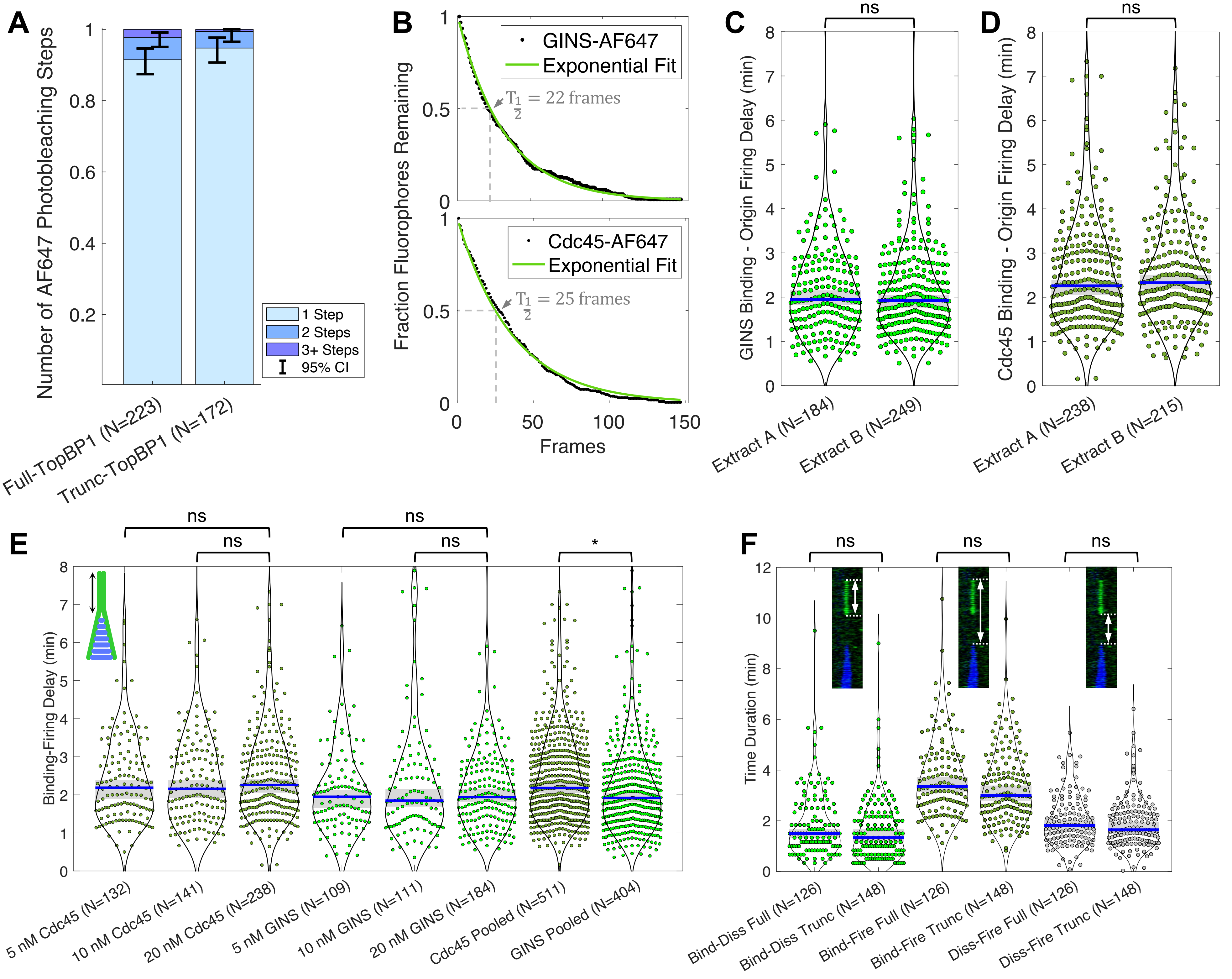


**Figure S7: Reproducibility and Technical Limitations, related to Figure 7**

**(A)** In vitro photobleaching analysis of full-length TopBP1$^{AF647}$ and truncated TopBP1$^{AF647}$ (error-bars – 95% CI). **(B)** Photobleaching decay curves for Cdc45$^{AF647}$ and GINS$^{AF647}$ and exponential fits to the data (with estimated half-lives indicated by $T_{1/2}$). Imaging conditions (exposure, laser power, TIRF angle) were identical to KEHRMIT experiments in this study. **(C)** Time delay between GINS binding and origin firing measured using two different egg extract preparations. **(D)** Time delay between Cdc45 binding and origin firing measured using two different egg extract preparations. **(E)** Time delay between Cdc45 or GINS binding and origin firing for 5 nM, 10 nM, and 20 nM concentrations of each protein. Experiments were conducted with the same extract preparation. **(F)** Properties of productive TopBP1 binding events (delay between TopBP1 binding and dissociation, delay between TopBP1 binding and origin firing, delay between TopBP1 dissociation and origin firing) compared between full-length TopBP1 and truncated TopBP1 constructs. **(C-F)** Blue bars and gray boxes represent the median and 95% CIs respectively.
